# Extracellular Vesicles of Salivary Mesenchymal Stem Cells Mitigate Acute Irradiation Injury: Use of an ex-vivo organotypic human slice tissue culture as a disease model

**DOI:** 10.1101/2025.09.04.674343

**Authors:** Akshaya Upadhyay, Migmar Tsamchoe, Anthony Zeitouni, Jordan Gigliotti, Junzheng Peng, Mahdokht Mahmoodi, Haider Abo Sharkh, Nicholas M. Makhoul, Michel El-Hakim, Jiangping Wu, Simon D. Tran

**Affiliations:** Faculty of Dental Medicine and Oral Health Sciences, Montréal, Québec, Canada, H3A 1G1; Research Institute of McGill University Health Center, Cancer Research Program, Montréal, Québec, Canada, H4A 0B1; Department of Anatomy and Cell Biology, McGill University, Montréal, Québec, Canada. H3A 2A7; McGill University Health Center, Royal Victoria Hospital, Montréal, Québec, Canada, H4A 3J1; Centre hospitalier de l’Université de Montréal, Montréal, Québec, Canada, H4A 0B1; Montreal General Hospital, Montréal, Québec, H3H 1V6

## Abstract

**Abstract:** Ionizing radiation (IR) therapy for cancer patients can damage surrounding healthy tissues, particularly the salivary glands (SGs), leading to oral and systemic health issues reducing the quality of life of the patients. The mechanisms behind IR damage in SGs are not fully understood, and current therapies often fail to meet patient needs adequately. Therefore, identifying targeted pathways and alternative treatments is essential.

To address this, we developed an ex vivo model of SG damage using human salivary glands obtained from patients. Healthy submandibular glands were harvested, cultured, and exposed to IR. RNA sequencing revealed elevated markers for DNA damage, inflammation, and ferroptosis, with four specific genes—FDXR, MDM2, H2AX, and p21—showing increases in expression that correlated with the IR dose. Using them, we developed a high-throughput genetic screening method to evaluate stem cell therapies aimed at mitigating IR injury.

Conditioned media from mesenchymal stem/stromal cells (MSC-CM) were found to reduce the expression of all four markers, maintain tissue viability, promote cell proliferation, and decrease oxidative stress. Further analysis involved separating MSC-CM into two fractions: Extracellular Vesicles (EV)-rich and EV-depleted. The EV-depleted fractions retained elevated levels of DNA damage response markers, indicating that EVs play a crucial role in mediating tissue repair. In contrast, the EV-rich fractions reduced the markers of DNA damage response and were readily absorbed by the tissue slices. In conclusion, we have developed a genetic screening method to evaluate treatments for acute IR injury, emphasizing the significant role that EVs play in the repair process.

## Background

Dry mouth, a sequela of iatrogenic irradiation damage to the salivary glands (SGs), increases the patient’s susceptibility to oral infections^1,2^, dental caries^3^, loss of taste^4^, and difficulty swallowing^5,6^, leading to considerable morbidity, malnutrition, and discomfort that can persist for several years^7,8^. Current therapies have limited efficacy and fail to meet patient satisfaction^9,10^. We have previously observed that Mesenchymal Stromal/Stem Cells (MSCs) reduce IR damage in murine models through paracrine secretions.^11,12^ We have further explored minor salivary gland-derived MSCs and their secretions to ameliorate IR damage in vivo.^13^ However, for the clinical translation of cell-based therapies, it is imperative to identify their active components and understand the repair mechanisms to develop targeted therapies that have minimal side effects.

SG research is challenging as the pre-clinical models replicating IR injury in humans are limited. Firstly, the saliva-secreting cells, acinar cells, are epithelial and more susceptible to stress and IR injury than other epithelial cells (namely, ductal cells) and mesenchymal cells.^14^ The only human-derived acinar cell line was developed in 1994, as maintaining epithelial cells in vitro cultures is challenging due to their propensity for epithelial-to-mesenchymal transition (EMT).^15^

Furthermore, the acinar-to-ductal transition and loss of polarization of acinar cells occur in stressful conditions.^16^ Secondly, SGs are composed of epithelial and mesenchymal components that are intricately intertwined. In addition to structural closeness, this also has a functional impact on their development^17^, homeostasis ^18–20^, and injury response^21^. IR injury and repair response is an amalgamation of vascular^22–25^, neural^26–30^, immune^29,31,32^, molecular^33–35^, or SG-specific pathways^20,36,37^. It becomes imperative to maintain these interactions in disease modeling for testing new therapeutics. While in vivo studies maintain these interactions, the studies using the current in-vivo models have a high bias, particularly in terms of performance and detection bias, as they often lack randomization and blinding^38^. Therefore, there is a need for robust molecular targets that can differentiate between different treatments, ideally with high precision, reproducibility, sensitivity, and specificity.

Here, we developed an organotypic tissue slice culture for IR injury using human Submandibular glands (SMGs). Slice culture is an ex vivo culture system comprising of tissues cut with precision into 100-150 μm slices and cultured over a porous membrane with culture media at the bottom (**Fig. 1a**). We aimed to maintain the tissue slices as closely as possible to their native conditions while ensuring hydration and nutrition. As no enzymatic digestion or chemical treatment was used to generate these tissue slices, the cell surface receptors and cellular connections remained intact^39^. Such ex vivo models effectively blend the efficiency of in vitro studies with the ability to analyze multiple test groups from a single donor. At the same time, they closely mimic the cellular and molecular environments found in in-vivo models. We used human SMGs, which are usually discarded during surgeries for benign tumor removal, mandibular reconstruction, or post-mortem.

**Fig1:**
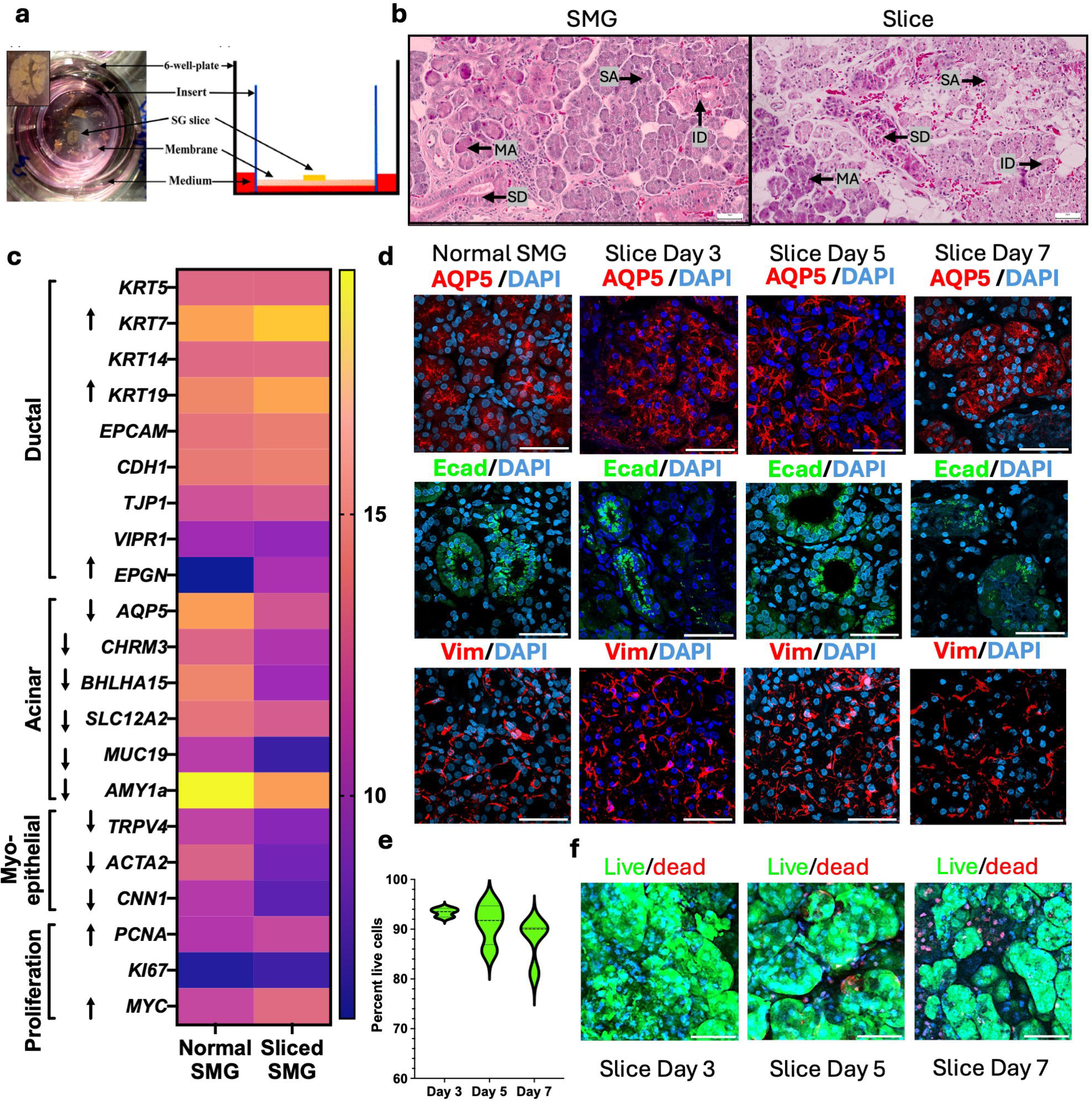
Human SMG organotypic tissue slice culture. **a** Experimental set-up for organotypic tissue slice culture (Adapted from Su et al., 2016)^51^. **b** Hematoxylin and Eosin staining (HnE) for normal SMG and SMG slices (SA=Serous acini, MA=Mucous acini, SD=Striated duct, ID=Intercalated duct, Scale bar = 50µm) **c** Immunofluorescence(IF) staining for SG markers-acinar cells: AQP5, epithelial cells: ECadherin, and mesenchymal/myoepithelial cells: Vimentin in normal SG and slices at day 3, 5 and 7, respectively(Scale bar= 50µm) **e/f** Live-dead assay for SMG slices at day 3, 5 and 7. Live cells are stained with Calcein, dead cells are stained with Ethidium Homodimer(ED1), and nuclei with Hoechst. The graph represents the percentage of live cells at each time point (n=3) (Scale bar= 50µm)

Furthermore, MSCs from various origins have been identified, which reduce IR damage in SGs^40–43^. We utilized labial SG-derived MSCs (LMSCs) for our study as have embryological proximity to SG development and comply with the FDA criteria for the use of a homologous source of stem cells as the target organ. Moreover, extracellular vehicles (EVs) derived from SG MSCs were more enriched in angiogenic proteins and showed comparable wound healing as adipose-derived and bone-derived EVs.^44^ SG MSC EVs were also more readily taken up by SG epithelial cells than Wharton’s jelly-derived MSCs.^45^

MSCs can be expanded from healthy donors, and their paracrine secretions can be utilized as either autologous or allogeneic treatments. This approach reduces the chances of immune rejection and enables the easy transport of secretions without a lag time in culturing cells from each patient. Additionally, the composition of these secretions can be assessed for quality, ensuring greater reproducibility and predictability compared to using whole cells. We isolated the MSC secretions, called Conditioned Medium (MSC-CM) by culturing them under serum free condition for 24 hours to induce the cells to secrete pro-angiogenic and anti-inflammatory factors.^46^ This process also increases extracellular vesicles (EV) secretions from the cells^47^.

In this study, we first identified the targets of acute irradiation damage in ex vivo salivary gland tissues from human donors using in-depth bulk RNA sequencing to create a high-throughput screen. We then proceeded to test the effect of MSC-derived Conditioned Media (MSC-CM) on IR injury using our ex-vivo model. Further, we purified MSC-CM to separate it into EV-rich and EV-depleted fractions, aiming to identify the biologically active component.

## 1. Results

### Organotypic tissue culture slices maintain heterogeneous cell types with salivary gland function in ex vivo culture

We compared the histology of human SMGs with that of sliced human SMGs using hematoxylin and eosin staining (**Fig. 1b**). The slices maintained the histological morphology, with all major salivary gland structures, namely acini (serous and mucous) and ductal structures (striated and intercalated), intact. Differential expression of genes was assessed using DeSeq2 to compare intact SMGs and SMG slices from the same four patients on day 2 (**Fig. 1c**). Selected ductal marker genes *KRT5*, *KRT14*, *EpCAM*, *CDH1*, *TJP1*, and *VIPR1* remained constant, while *KRT7*, *KRT19* and *EPGN* were upregulated (log2FC >2, p-adj < 0.05). Additionally, proliferation markers *PCNA* and *MYC* were upregulated in SMG slices (log2FC > 2, p-adj < 0.05). Acinar cell markers, *AQP5*, *CHRM3*, *BHLHA*15, *SLC12A2*, *MUC19*, *AMY1A*, *ASCL2*, *ASCL3*, and myoepithelial markers *TRPV4*, *ACTA2*, *CNN1* were all downregulated (log2FC >2, p-adj < 0.05). However, the transcript counts remained above the detection limits. Log fold change with the counts and p-adjusted values for the selected SG, endothelial, and macrophage markers are provided in Supplementary Table **T4**. To test if this change in gene expression affected the expression of the proteins, we tested for E-Cadherin (ductal), AQP5 (acinar water channel), and Vimentin (mesenchymal and myoepithelial (MECs)) using Immunofluorescence (IF) staining over the day 2, 4 and 6 of culture (**Fig 1d**). The intensity of these markers, normalized to intact tissue (**Fig S1a**), remained the same for AQP5, decreased for E-cadherin, and increased slightly for Vimentin on day 3. This was expected, as mesenchymal/MECs exhibit a reparative response in vitro^48^; there may also be some degree of EMT occurring for the purpose of cell survival. A slight elevation in the expression of all three was observed on day 5. However, by day 7, all the markers decreased in expression. Furthermore, AQP5 is a water-channel protein essential for the secretion of saliva, expressed at the basolateral surface of acinar cells, which facilitates the secretion of saliva into the lumen. Its expression was maintained and localized at the basolateral side of the acinar cells, giving a characteristic spider web appearance. This is a significant finding for us as pre-clinical salivary gland models often lose AQP5 polarization, which is crucial for acinar cell function. Amylase is a predominant protein present in saliva; its expression was also confirmed by western blot staining (Data not shown). Over 6 days, the survival of tissue culture slices was assessed using a live-dead assay, where live cells were stained with Calcein, dead cells with Ethidium Homodimer (ED-1), and nuclei with Hoechst (**Fig. 1e and f**). The normalized live cell count was 93.4±0.9%, 91.1±4.1%, and 87.9±4.5% for days 3, 5, and 7, respectively.

### Organotypic tissue culture is a model for ex vivo IR injury in SMGs

We opted for a 3-day protocol to maintain optimal SG protein expression, function, and viability (**Fig. 2a**). Tissue slices from fresh SMGs were cut and cultured on day 1, followed by IR exposure and treatment on day 2 (24 hours post-IR) and sample collection was done after 12-24 hours on day 3. To identify the key genes responsible for IR damage, we performed bulk RNA sequencing followed by differential count analysis using DeSeq2 to compare gene expression in IR-exposed SG slices with those in control (no IR) slices 24 hours post-IR. Various DNA damage (*MDM2*, *H2BC4*, *H2BC21*), apoptotic (*CDKN1A/p21*, *TIGAR*), ferroptotic (*FDXR*) and inflammatory response genes (*VWCE*, *TNFRSF10C*, *Ly6D*) were significantly upregulated (**Fig 2b**). Gene enrichment using Metascape identified pathways related to p53, DNA damage, and cellular response to stress as being upregulated. At the same time, the cell cycle and DNA metabolic processes were downregulated (**Fig.S1b,c**), which is consistent with previous in vitro studies.^49^ To determine the targets for IR damage, we exposed the tissue slices to various IR doses. Linear regression analysis for the expression *of MDM2*, *FDXR*, *H2AX,* and *p21* in response to different IR doses showed R^2^ values close to 1, showing linear goodness-of-fit of these genes, with a maximum for *p21*(R^2^=0.9795), followed by g*H2AX*(R^2^=0.9388), *FDXR*(R^2^=0.9263) and *MDM2*(R^2^=0.9201). All of them yielded significant p-values (≤0.05); therefore, we selected these four genes as a screen to identify IR damage in various samples for further experiments. Furthermore, gamma-H2AX is a phosphorylated form of H2AX, which is expressed at the breaks in the double-stranded DNA in response to damage, particularly IR damage. We observed a progressive increase in the immunofluorescence of gH2AX foci in response to IR (**Fig S1d**).

**Fig2:**
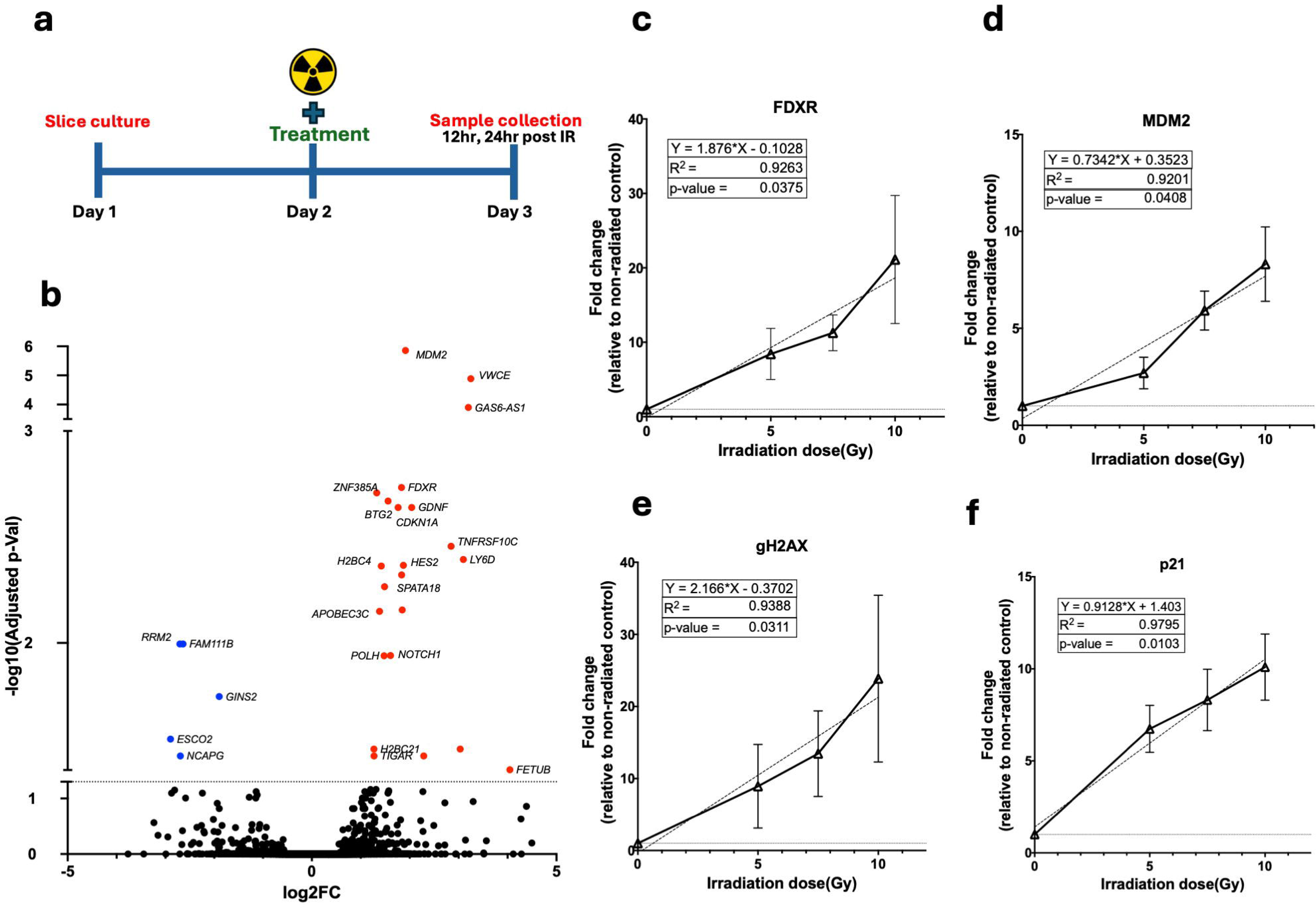
Human SMG Slice culture as a model to study radiation injury. **a** Schematic image of time points of culture, IR exposure/treatment and sample collection. **b** Volcano plot of differentially expressed genes (DEGs) in response to IR in sliced tissue homogenates when compared to non-IR control slices (n=4, p-Adj < 0.05, log2FC >2) **c-f** qPCR for selected DEGs in response to 5Gy, 7.5GY, and 10Gy IR dose. Fold-change in control (with no IR) is normalized to 1. MDM2, FDXR, H2AX and p21 show a dose dependent IR response in slices after 24 hours of IR exposure(n=3).

### MSC conditioned media (MSC-CM) reduced IR-induced damage in the tissue culture slices

To identify the active components in MSC secretions, we collected the conditioned media derived from serum-free starved MSCs (MSC-CM). The optimal protein concentration of MSC-CM for treatment was identified using viability tests in in-vitro SMG cell culture (**Fig S2a**). A concentration of 25 μg/ml for 2 hours post-IR exposure increased viability and restored proliferation in IR-exposed SMG cells. The control group was taken to the irradiator room but not exposed to IR or MSC-CM treatment. The IR group was exposed to 7.5 Gy of IR, followed by the addition of PBS only. The IR+CM group was exposed to IR, followed by the addition of MSC-CM.

Using the previously identified genes, we performed quantitative PCR (qPCR) on mRNAs extracted from sliced tissue lysates 24 hours after IR exposure (Day 3 of culture). As expected, we observed a significant increase in *MDM2*(p≤0.001), *FDXR*(p≤0.001), *H2AX*(p≤0.01), and *p21* (p≤0.01) expression after IR exposure (**Fig 3a**). Their expression was reduced in response to IR with MSC-CM treatment, and a significant change was observed in *H2AX* only (p ≤ 0.01). The increase in *MDM2*, *FDXR*, and *p21* expression was less significant after MSC-CM treatment than after no treatment following IR exposure, compared to the control. We validated the expression of gH2AX using immunofluorescence staining to visualize double stranded DNA break directly (**Fig 3b, c**). Using particle count analysis (normalized to the number of nuclei), there was an increase in the expression of gH2AX positive nuclei in the IR-only group (p≤0.05). This increase was significantly reduced after treatment (p≤0.05) to the level of the control group. Furthermore, post-24-hour IR, there was a variable decrease in the percentage of live viable cells (p ≤ 0.05) (**Fig. 3d, e**). In slices with IR and MSC-CM treatment, the viability was significantly higher than in the IR-only group (p ≤ 0.01). Proliferation in the tissues was also assessed by Western blot of the tissue lysates and staining for Ki-67; however, no significant difference was observed (data not shown). Moreover, the tissue culture supernatant was collected to assess the release of superoxide dismutase enzyme, which is secreted in response to oxidative stress. A slight increase in SOD activity in the tissue culture media (SOD3) was observed after IR (p ≤ 0.01). The SOD activity was less elevated in response to treatment (p≤0.05) (**Fig S2b**).

**Fig3:**
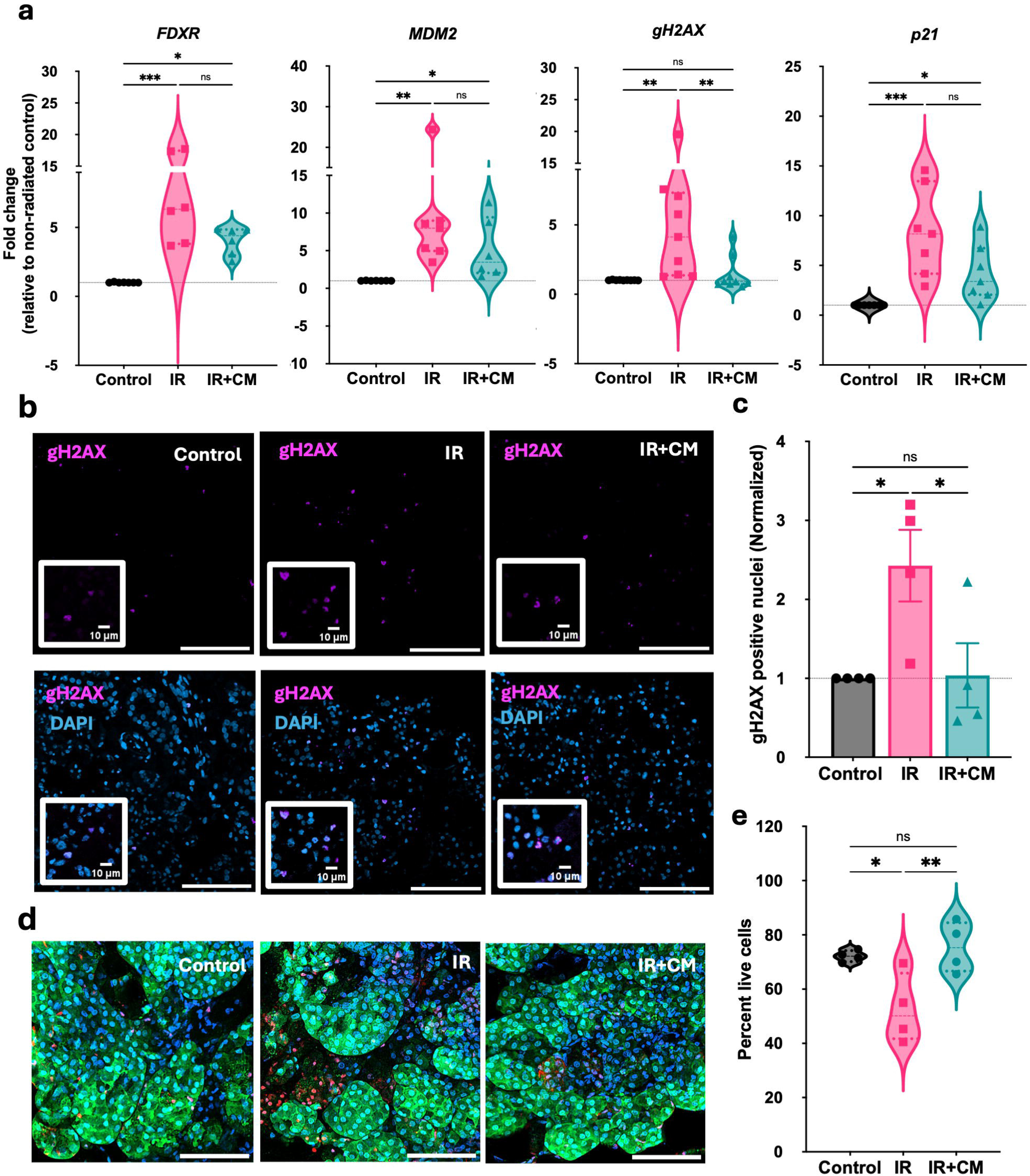
Human MSC CM treatment mitigates IR-induced injury in the tissue culture slices model. **a** qPCR fold-change in tissue lysates for IR response genes *MDM2*, *FDXR*, *gH2AX*, and *p21* in control (no radiation, no treatment), IR (irradiated) and IR+ Treated (irradiated and treated with MSC-CM) SG slices (Kruskal-Wallis test with Dunns multiple comparison) **b/c** Immunofluorescence staining γH2AX (magenta) with DAPI(cyan) in tissue culture slices with mage analysis for γH2AX particle count using ImageJ, normalized to control(n=5, One-way ANOVA with Tukey’s multiple comparison test, scale bar=100µm) **d/e** Live-dead staining of viable tissue culture slices after 24 hours of IR with particle count analysis for number of ED-1 positive nuclei normalized to Hoechst using ImageJ software (One-way ANOVA with Tukey’s multiple comparison test, scale bar= 50µm)

### Extracellular vesicles (EV) depletion from MSC-CM reduced its bioactivity

We further tested various fractions of MSC-CM to identify the bioactive components of MSC-CM: whole CM, EV-depleted CM (soluble fraction), and EV-rich CM (vesicular fraction). From here onwards, the tissue slices group treated with MSC-conditioned media is labeled as IR+CM, the EV fraction is IR+EV, and the soluble fraction is IR+EV-depleted. We further added two additional groups, in which heparin sulfate (10μg/ml) was added along with the EV fraction to inhibit the uptake of EVs, and the EV membrane was lysed with KCl to identify the role of the membrane in mediating the biological response^50^.

Transmission Electron Microscopy (TEM) imaging of EVs revealed intact, double-membrane spherical structures that identified EVs (**Fig. 4a**). Markers for EVs were characterized using Western blot with complete cell lysate from MSCs used as a control. The EV-rich fraction was positive for CD63 (not shown), TSG101, and Syntenin and negative for Calnexin (**Fig 4b**), while the EV-depleted fraction lacked CD63 (not shown) and Syntenin. Therefore, it was confirmed that the EV isolation was complete, and the EV-depleted fraction was completely deprived of EVs. 8.7e10 ± 3.5e9 particles/ml were observed with an average size of 105 nm using Nanoparticle tracking analysis (NTA), which indicates the presence of pure EVs (size= 50-150 nm) (**Fig 4d**). MSC-CM consisted of various growth factors as identified through Mass Spectrometry (data not shown). As these growth factors can be involved in IR repair, we wanted to see how many of these are carried over in the EV and EV-depleted fractions. We performed Human Angiogenesis & Growth Factor 17-Plex Assay to test presence of Angiopoietin-2, BMP-9, EGF, Endoglin, Endothelin-1, FGF-1, FGF-2, Follistatin, G-CSF/CSF-3, HB-EGF, HGF, IL-8/CXCL8, Leptin, PLGF, VEGF-A, VEGF-C and VEGF-D. Most growth factors were conserved in the MSC-CM and EV-depleted fraction, as expected since they are commonly found in the soluble secretions of cells (Fig **S2c**). Expression values for BMP-9, Endothllin-1, FGF1, HbEGF, and VEGF-D were below 10, so they were excluded to avoid extrapolation of the results. To identify the growth factors carried over in EVs, we normalized the observed concentrations using the Endoglin concentration in CM and the EV fraction (**Fig. 4c**). This is because endoglin (a.k.a. CD105) is a membrane protein in EVs, which was enriched in EV preparations as expected. However, their high concentration in the EV fraction is an indicator of the large number of EVs that had to be lysed to obtain the same number of proteins for the assay, which required a standardized protein concentration. Growth factors such as FGF2, VEGF, and leptin were detected at approximately 10-15% in the EV fractions, while traces of EGF, FGF1, and VEGF-A were also present. Whether these factors are carried inside the EVs or are co-isolated will have to be studied with further investigations. EV uptake was confirmed by SMG cells, where EVs were internalized by the cells, as observed by PKH67 labeling (**Fig. 4e**).

**Fig4:**
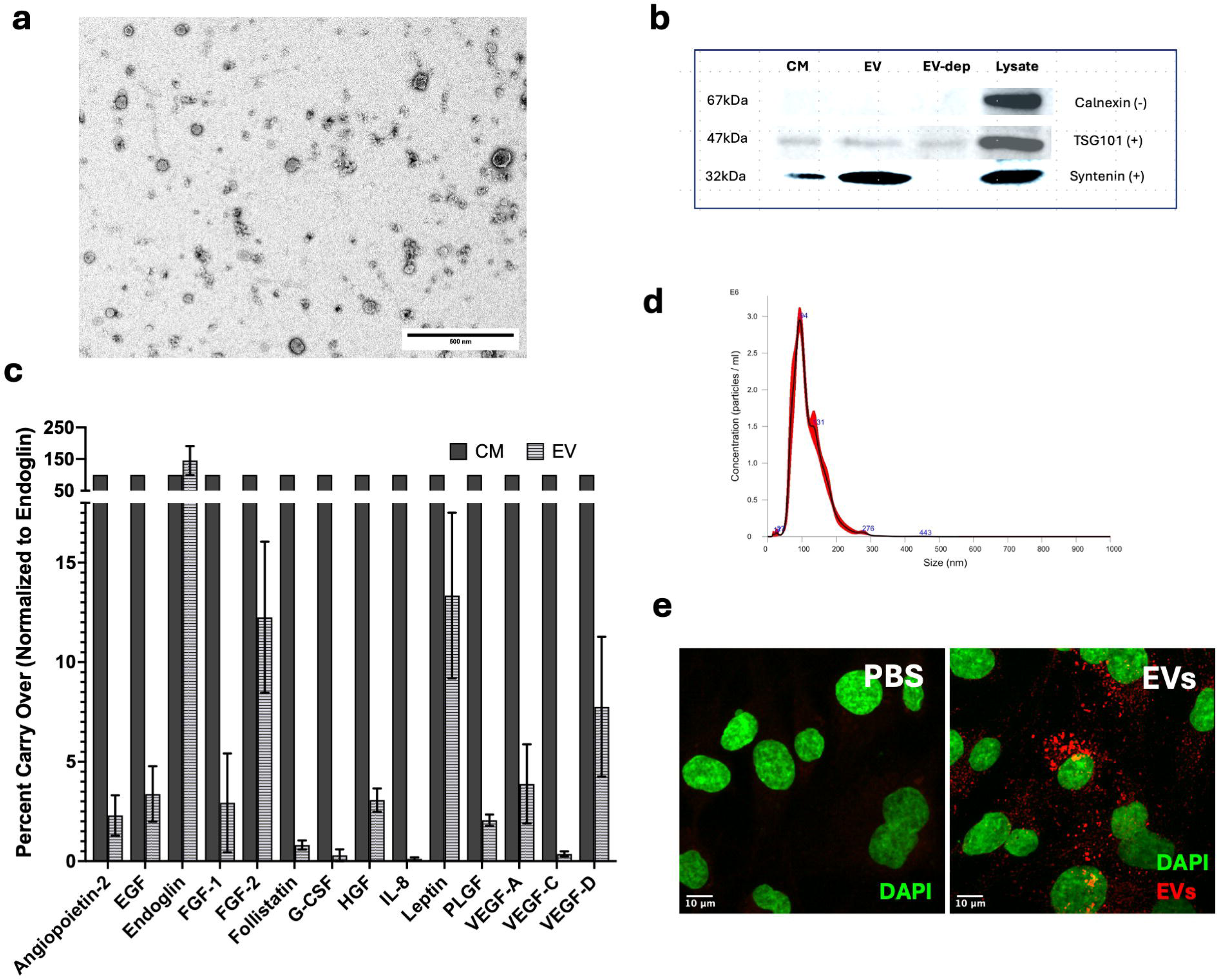
EV characterization. a TEM image of MSC-EVs negative staining under 13000X b Western blot of Conditioned media (CM), extracellular vesicle (EV), EV-depleted fraction and whole cell lysate, stained for EV-specific markers. c Multiplex analysis of human angiogenesis growth factors present in CM and EV fractions normalized to Endoglin levels in CM and EV respectively* d Size distribution and count for MSC-EV using Nanoparticle Tracking Analysis(NTA) f Pkh67 labeled filtered-PBS control and EVs in human SMG cells (Scale bar=10µm) *BMP-9, FGF-2, Hb-EGF and endothelin-1 are not shown as the expression levels we low (0-1.2pg/ml). EGF= Epithelial growth factor, FGF= Fibroblast growth factor, G-CSF= granulocyte-colony stimulating factor, HGF= Hepatocyte growth factor, IL= Interleukin, PLGF= Platelet derived growth factor, VEGF= Vascular endothelial growth factor

Using our identified genetic screen, we then proceeded to test these various fractions of CM in IR-damaged tissue culture slices. There was an increase in *MDM2*(p≤0.05), *FDXR*(p≤0.01), *H2AX* (ns) and *p21*(p≤0.01) expression at mRNA level 24 hours after IR. After treatment with the EV-depleted fraction, the expression of *FDXR*, *MDM2*, and *p21* was sustained over 24 hours. Thus, DNA damage and ferroptosis were significantly high (p≤0.01, p≤0.05, and p≤0.01, respectively). To assess the translation of these proteins, we tested the expression of 7 DNA damage proteins 12 hours and 24 hours after IR (**Fig 5b, S2d-h**). We chose the 7-plex assay because it contained most of the genes identified in our previous genetic screen, namely ATR, Chk1 (Ser345), Chk2 (Thr68), H2A.X (Ser139), p53 (Ser15), MDM2, and p21. MDM2 was significantly elevated at 12 hours post-IR, while p21 remained elevated at 12- and 24-hour post-IR. p21 also remained elevated in the EV-depleted group and the EV-heparin sulfate combination. However, it is worth noting that Heparin Sulfate reduced the panel markers at the mRNA level. Furthermore, the removal of the EV membrane using KCl completely lost the bioactivity of the EV fraction as the DNA damage proteins remained high (data not shown).

**Fig5.**
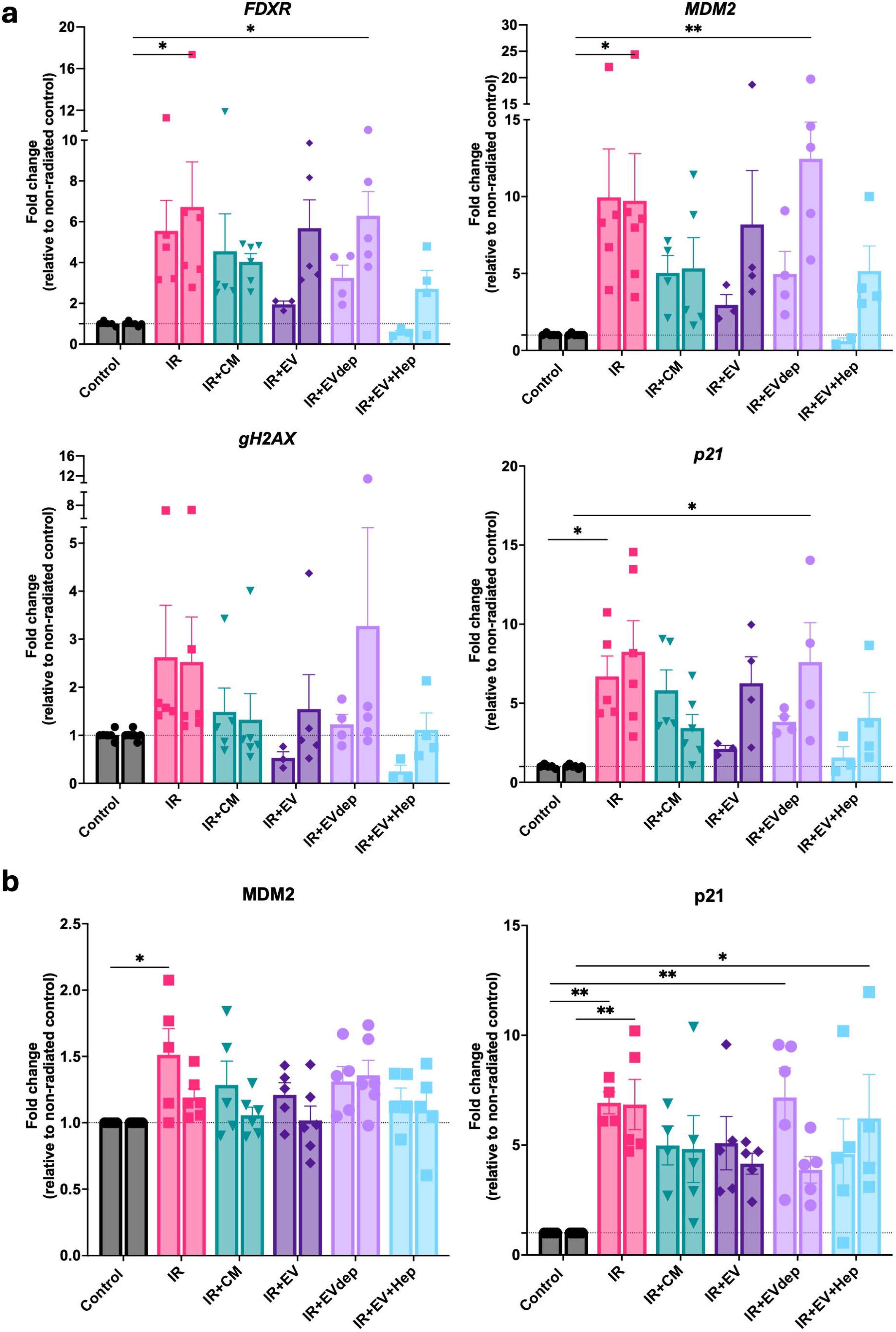
IR response after different MSC-CM fraction treatment. a qPCR for FDXR, MDM2, gH2AX and p21 normalized to Control at 12 and 24-hour post-IR, respectively (Two-way ANOVA with Tukey’s Multiple comparison test) b DNA damage response protein multiplex assay in the tissue homogenates 12-hour and 24-hour post-IR, respectively. Fluorescence intensities were normalized to the control values to obtain fold-change in the protein concentrations (Two-way ANOVA with Tukey’s Multiple comparison test for each protein expression at 12 and 24 hour)

## 2. Discussion

### Ex-vivo organotypic tissue culture slice culture as a disease model for salivary glands and acute IR injury

Previously, the slice culture model was observed to have intact acinar cell clusters, ductal cell clusters, nerve innervation, and small vasculature. ^39^ The slices also maintained the physiological role of the glands by maintaining the ability to transfer Calcium ions across channels in response to sympathetic and parasympathetic stimulation, which is required for saliva secretion.^39^We also observed that various salivary gland markers crucial for salivary gland (SG) function were expressed and localized, similar to those in normal SGs. There was a gradual decrease in both epithelial and mesenchymal markers, consistent with previous findings from our group.^51^ It could be attributed to the stress during processing and prolonged culture of the tissues. To maintain optimal conditions, we proceeded with a 3-day protocol.

For natural extracts and derivatives, high- to moderate-throughput screens are required to compare bioactivity. To our knowledge, this is the first attempt to comprehensively identify acute response genes in ex vivo human SGs using RNA sequencing and systematically validate target genes with different dose responses using qPCR. Previously, doses of 0-15 Gy have been tested in in vitro SG culture models^52–57^. Following these studies, we also tested doses of 5, 7.5, and 10 Gy. 7.5 Gy, which is approximately the average cumulative clinical dose of a week, was chosen for the subsequent treatment experiments to get the maximum IR damage output without complete tissue death^52,56^.

Among all the upregulated genes, the DNA damage response genes *MDM2*, *gH2AX*, and *p21*, as well as the ferroptosis gene *FDXR*, showed a dose-dependent response to IR exposure. These findings are consistent with the dosimetry studies performed with blood samples for triage of IR-exposed individuals^58–61^. MDM2, gH2AX, and p21 are well-known DNA damage response proteins downstream of ATM and ATR. Recently, ATM has also been shown to be essential for ferroptosis^62,63^. Our findings show that in addition to apoptosis, ferroptosis occurs in response to DNA damage in SGs. Interestingly, the glial-cell-derived neurotrophic growth factor (GDNF) was also upregulated in our samples exposed to IR, which corresponds with previous observations from clinically IR-exposed patient biopsy samples.^48^ The role of GDNF was also previously identified in SG survival post-IR, primarily mediated through MECs.^64^ As our culture maintained the MEC population, we can extrapolate to conclude that this crucial interaction is maintained in our culture.

H2AX is activated through phosphorylation at serine 139^65^(then called gH2AX) by various IR response genes, principally ATM^66^. gH2AX expression was increased after IR in clinically radiated human SMG samples 3-4 months after IR and mice after 4-48 hours post-IR^67^. In another study, it increased within 30 minutes to 1 hour but not at 24 hours.^68^ We observed the increased expression of gH2AX at 12 and 24 hours in a dose dependent manner, consistent with previous studies^69^.

These findings are significant as this model can be used efficiently to compare different therapies on the same patient donor. It will ensure the reproducibility and reliability of biological and drug therapies. It also offers advantage of studying the effect of IR in various patient samples closer to their native state, especially older individuals where there are higher chances of fibrosis limiting cell and organoid cultures. However, a limitation of our model is that it does not accurately represent chronic IR injury, which can involve inflammation and fibrosis. The long-term effects of these acute IR response markers are yet to be elucidated using in vivo models.

### MSC-CM can reduce DNA damage in ex-vivo SGs through EVs

Conditioned media is a biologically complex secretion from MSCs obtained after concentrating serum-free MSC culture media. All four identified IR response markers were reduced after treatment with MSC-CM. PTGS2 and CHAC1 are additional markers of ferroptosis^70^; we also observed their reduction at the mRNA level (data not shown). Thus, we can conclude that both DNA damage and ferroptosis were reduced after MSC-CM treatment. The live-dead assay provides a real-time image of tissue viability. Live cell areas were drastically reduced in response to IR but restored upon treatment in our study, which is consistent with the findings of Shin et al^71^. Song et al. did not observe significant changes in live-dead cells after IR and treatment 2 days post-IR, possibly due to the high IR dose used (15 Gy)^69^ . MSC-CM maintained the viability of the tissue slices, promoted proliferation in irradiated primary salivary gland cells, and reduced oxidative stress.

Furthermore, conditioned media consists of vesicular and soluble fractions, which can be separated through various methods. We chose differential ultracentrifugation to separate them. It enabled us to separate the EVs from the soluble fraction (EV-depleted) without the addition of any other reagents or chemicals so we could also use the EV-depleted fraction for comparison.

The soluble components consist of growth factors and cytokines that are released into the extracellular environment. On the other hand, the vesicular fraction is characterized by a range of small membrane-bound structures, including apoptotic bodies, microvesicles (MVs), and EVs. EVs have garnered increasing interest among researchers because they are generated through a sophisticated endosomal pathway, which allows them to facilitate precise communication between cells. EVs from embryonic mesenchyme interacted with epithelial cells and were essential for SG development^17^. In our study, after treatment with EV-depleted fractions (soluble component only), the IR response genes and proteins were significantly elevated after 24 hours, indicating a prolonged effect of IR remaining in these samples. EV-rich fractions reduced DNA damage response marker gene expression at both 12 and 24 hours and were internalized by the SMG cells, consistent with the findings of Kim et al^45^ and Su et al^40^. However, at the protein level, the difference was not apparent, which could indicate a higher dose of EVs is required. Additionally, treatment with Heparin Sulphate before MSC-CM was done to block the EV and cellular interaction^50,72^. However, we observed a further reduction in our target markers. This anomaly can be explained as Heparin Sulphate itself can affect the repair process, thus acting synergistically with the EVs.^73^

We found traces of some growth factors in the EV fractions. EVs can carry various growth factors. For instance, HGF^74^ can also be packed and transferred^75^ into EVs, as can multiple other proteins^76,77^. Moreover, EVs can be required to protect and uptake active factors^78^. Whether these factors are carried inside the EVs or are co-isolated will have to be studied with further investigations. SG MSC EVs were observed to be enriched in miRNAs that suppress hypoxia-induced autophagy and anti-inflammatory pathways.^45^ Given that the amount of growth factors is low in the EV-rich fractions, miRNA composition can also be studied to identify active factors present in the EVs. Future studies will investigate the protein and miRNA composition of EVs to identify the active factors. It will aid in the clinical translation of MSC therapeutics, ensuring minimal side effects.

## 3. Methods

### EXPERIMENTAL MODEL AND SUBJECT DETAILS

#### Human samples

The study was approved by the McGill Institutional Review Board (IRB A05-M62-05B). From 2022-2024, freshly resected healthy Submandibular glands were obtained from 18 adult patient donors, both male and female, aged 18-80 (51.3 ± 21.4 years) undergoing oral, maxillofacial and ENT surgeries due to benign conditions, fracture, or post-death from Royal Victoria Hospital, Montreal General Hospital, and Transplant Quebec. The demographics of the patients are provided in Supplementary Table **1**.

#### Organotypic tissue culture and irradiation

Tissue culture slices were generated using our previously established protocol^79^. Briefly, the glands were rinsed in sterile Phosphate-Buffered Saline (PBS), connective tissue was removed, and they were cut into 3-4 mm size pieces, then embedded in a 3% low-melting agarose gel (Sigma Aldrich, A9414-50G). The gel-covered tissues were glued on the platform of the vibratome(Leica BiosystemsVT1200), cut into viable 100-150μm thin tissue slices in ice-cold PBS, transferred in Hanks Balanced Salt Solution(Gibco, 14025092) at room temperature, washed twice with 2% anti-anti PBS and spread over Trans well membrane (Millipore Sigma, PICM0RG50) with Mammary Epithelial Basal Medium (EpiMax1) (Wisent Inc., 002-010-CL) with 5% Fetal Bovine Serum(FBS) below the membrane in a humidified cell culture incubator (37°C, 5% CO_2_).

The RS 2000 X-ray Biological Irradiator, available at the Comparative Medicine and Animal Resources Centre, McGill University, was used for IR exposure. The time of exposure for each dose was calculated according to the calibration of the irradiator. SG slices were cultured over 24 hours, followed by IR with appropriate doses (and treatment) and sample collection 12–24 hours post-IR.

### METHOD DETAILS

#### Conditioned media preparation and EV isolation

LMSCs were cultured in Minimal Essential Medium (MEM, Gibco 12561056) supplemented with 10% FBS and 1% antibiotics in 150mm tissue culture-treated dishes until 70-80% confluency was reached. MSCs were then washed twice with PBS and cultured in serum-free conditions for 24 hours. MSC viability was assessed after each starvation (TC20 Cell Counter, Bio-Rad 1450102) to ensure that cell viability was above 90%. The culture media, now referred to as conditioned media (CM), was collected and centrifuged at 500g for 5 minutes at 4°C, followed by centrifugation at 3000g for 20 minutes at 4°C to remove cell debris. The CM was then concentrated using an Amicon tube with the required pore size (3 kDa/10 kDa/100 kDa, Millipore Sigma Amicon Ultra-15 Centrifugal Filter) from 15 mL to 150 μL (3500 RPM for 30 minutes at 4°C). MSC-CM protein concentration was assessed using the MicroBCA Protein Assay Kit (Thermo Scientific, 23235) according to the manufacturer’s instructions and was stored at -80 °C until further use.

For EV characterization, we followed MISEV guideline ^80^. CM was reduced from 100 mL to 1 mL using Amicon 10 kDa filter tubes and filtered through a 0.22 μm filter. CM was kept on ice and transferred as required during all procedures. 500 ml of Concentrated CM was stored for the CM treatment group, and another 500 μl was subjected to ultracentrifugation at 110,000 × g for 2 hours at 4°C (MLA 130 rotor, Beckman Coulter OptimaMax-130k Tabletop Ultracentrifuge)^72^. The EV-depleted supernatant was collected for the EV-depleted fraction, while the pellet was further washed at 110,000×g for 2 hours at 4°C in filtered PBS and then suspended in 500 μL filtered PBS for the EV-rich fraction. The supernatant over the pellet collected from the first ultracentrifugation was also subjected to ultracentrifugation at 110,000 × g again to ensure the CM was completely devoid of EVs. The CM, EVs, and EV-depleted CM were aliquoted into 50 mL fractions to prevent freeze-thawing and stored at -80 °C for up to 3-4 months.

For treatment, 25 μg/mL CM (or EV-depleted CM) was added to the EpiMax1 culture media below the Trans-well membranes immediately after IR exposure and replaced with only EpiMax1 media after 2 hours of incubation. EV treatment was normalized based on volume of MSC-CM. 10μg/ml Heparin sulfate was added 2 hours before EV treatment to inhibit EV uptake^72^. For EV membrane removal, 200 μL of EVs was mixed with 200 μL of 2 M KCl solution for 30 min at 4°C with agitation. The preparations were then subjected to ultracentrifugation at 110,000 × g for 1 hour to pellet the treated EVs. ^72^

#### Immunofluorescence and staining

The tissue slices were stained using our previously established protocol^79^. Briefly, we fixed the tissue culture slices in 4% paraformaldehyde overnight at 4°C, processed and embedded them in paraffin, and cut them into 5μm sections. For dewaxing, the slides were heated at 95°C and incubated in CitriSolv solution for 5 minutes twice, followed by rehydration in sequentially decreasing concentrations of ethanol. Antigen removal was done using Sodium Citrate buffer at 95°C for 30 minutes, permeabilized using 1% Triton for 5 minutes, blocked with Universal blocking solution(Biogenex, HK085-5K), primary antibodies (Supplementary table **T2**) were added in appropriate concentrations overnight at 4°C in a humidified chamber, followed by secondary antibodies (Alexa Flour 594 anti-Rabbit, FITC anti-mouse, Thermo Fischer) and 4, 6-diamidino-2-phenylindole, dihydrochloride (DAPI) staining after PBS-Tween 20 wash(ThermoScientific, 85113). Slides were washed and mounted with Dako fluorescent mounting media(S3023). Stained slides were visualized using a Confocal microscope (Zeiss LSM 900) at 20X and 40X oil immersion with the appropriate lasers. At least 4-5 areas were imaged at 20X for analysis with 2-3 different slices for each group of each patient.

#### Live-dead assay and confocal microscopy

The tissue slices were stained with a Live/Dead viability/cytotoxicity assay kit (L3224, Life Technologies, Thermofischer). 2μl/ml Calcein AM, and 4μl/ml of EthD1 in PBS with Hoechst 33342 (1/20,000) (Invitrogen, USA) was used. Tissues were washed twice for 5 minutes each with PBS, then incubated with 100-200 μL of combined live/dead assay reagents for 1 hour at 37°C. The tissues were washed twice with PBS for 5 minutes each, followed by imaging under a confocal microscope (Leica LSM900). Images were taken with low magnification(10X) at multiple positions to get a broader area for image analysis with Z-stacks (5-10μm thickness) acquired and reconstructed using a maximum intensity tool to create the 2D images in Fiji ImageJ software. Dead cell counts were measured using particle count on Fiji ImageJ software and normalized against the number of nuclei stained with Hoechst (blue).

#### RNA isolation and qPCR

RNA and protein isolation for tissue culture slices were done simultaneously, as previously described^79^. Briefly, tissue slices were collected at the required time points and stored in RNAlater solution (Invitrogen, AM7021) at -80°C. RNA and protein were isolated using the mirVana PARIS kit (AM1556) according to the manufacturer’s instructions and stored in -80 for further analysis. Total RNA was quantified using a Nanodrop (Thermo Fisher Scientific, ND-2000). cDNA was prepared using the Revert Aid First Strand cDNA Synthesis Kit (Thermo Fischer, K1622) according to the instructions. For quantitative PCR, we used PowerUp SYBR Green Master Mix (Applied Biosystems, A25742) and primers (Supplementary Table **T3**) with 4-8μg cDNA in 10μl of the mix using the default long SYBR Green cycle on Step-One plus real-time PCR system (Applied Bioscience). The relative expression of the genes of interest was normalized to GAPDH^51^, and ΔΔCt values were calculated^81^. Three technical replicates were tested for each sample.

#### RNA sequencing and bioinformatic analysis

For RNA-Seq, the RNA samples were sent to Genome Quebec in Montreal, Quebec. mRNA-stranded libraries were prepared after a quality check. Sequencing was performed with NovaSeq6000 S4 PE 100B (Illumina) at a depth of 50 million reads, following the manufacturer’s protocol. The data was uploaded on Cedar servers provided by Digital Alliance of Canada for further analysis. A quality check for the read files was performed using MultiQC; no trimming was required. The reads were then mapped against the Grch38 (Genome Reference Consortium Human Build 38) human genome index using the STAR aligner, quantified using featureCounts, and significant differentially expressed genes were identified through the DESeq2 pipeline with a threshold of p < 0.05, and a fold change larger than 2 using RStudio. Functional annotation to identify core pathways was performed using the Metascape database (http://www.metascape.org) (Minimum overlap ≥ 3 and p < 0.05).

#### DNA damage multiplex assay

Luminex xMAP technology was used to quantitatively and simultaneously detect seven DNA damage and repair proteins. The multiplexing analysis was performed by Eve Technologies Corporation (Calgary, Alberta, Canada) using the Luminex® 200™ system (Luminex Corporation/DiaSorin, Saluggia, Italy) with Bio-Plex Manager™ software (Bio-Rad Laboratories Inc., Hercules, California, USA). Seven markers were measured in the samples using the Eve Technologies’ 7-plex DNA Damage/Genotoxicity Custom Assay as per the manufacturer’s instructions for use (MILLIPLEX® 7-Plex DNA Damage/Genotoxicity Magnetic Bead Kit Cat. # 48-621MAG, MilliporeSigma, Burlington, Massachusetts, USA). The 7-plex consisted of ATR, Chk1 (Ser345), Chk2 (Thr68), H2A.X (Ser139), p53 (Ser15), MDM2, and p21. Additional information can be found in the MilliporeSigma MILLIPLEX® protocol 48-621MAG.

#### NTA for MSC-EVs

The size and number of EVs were determined using a NanoSight NS300 from Malvern Panalytical. EV preparations were diluted to 1/100-1/500 to achieve 40-80 particles per frame. EV-depleted and filtered PBS were used as the negative control. Four 60-second videos were obtained for each sample at random time points automatically, with the viscosity set to water, the camera level set to 16, and under automatic detection. The threshold was set to 8, and the blur size was set to automatic at a temperature of 23.5-23.6°C. 8.7e+10 ± 3.5e+9 particles/ml were observed with an average size of 105nm. Each 1 mL of EVs was extracted from ten 150 mm round cell culture dishes, with an average of 3 × 10^6 cells in each.

#### TEM and confocal for MSC-EVs

EVs suspended in PBS were fixed into carbon-coated grids using glutaraldehyde. EV visualization was performed using negative staining with uranyl acetate and observed under a FEI Tecnai G2 Spirit Twin 120 kV Cryo-TEM at the Facility for Electron Microscopy Research, McGill University. Images were obtained with appropriate magnification.

For PKH67 labeling, EVs from concentrated CM were passed through an Izon column (IZON, IC135) to separate EVs from soluble proteins, thereby preventing multiple ultracentrifugation cycles. EVs were labeled using the manufacturer’s protocol. Briefly, diluent C and PKH67 dye were added in the required amounts, and the mixture was incubated in the dark at room temperature for 30 minutes. The reaction was then quenched by adding 10% BSA. The labeled EV particles were obtained in a pellet by sucrose gradient ultracentrifugation at 110,000 × g for 1 hour (Ti70 rotor, Beckman Coulter Optima L-100XP Ultracentrifuge) and concentrated in Amicon tubes (10 kDa filter). EVs were added to the slices and SMG cells for two hours, washed with filtered PBS 3 × 5 minutes, fixed with 4% PFA, and visualized with DAPI staining for nuclei, imaged under the confocal microscope (Zeiss LSM900, 63Xoil) by adding a drop of mounting media on the slide and covering with a cover slip.

#### Western blot and proteomic analysis for MSC-EVs

Proteins were extracted from MSCs for cell lysate control using a mirVana PARIS kit (AM1556). For EVs, 10% 10X RIPA (AB156034-1001) with protease inhibitor (Millipore, 535142) was added for 30 minutes on ice. Samples were prepared using Laemmli Buffer and β-mercaptoethanol (9:1), a 3-part buffer, and 1 part sample. 7-10μg of protein from each was added to 10% dodecyl sulfate-polyacrylamide gel electrophoresis (SDS-PAG) gel. After resolving the proteins sufficiently, the gels were transferred to a PVDF membrane, blocked with 5% milk for 1 hour at room temperature, incubated with primary antibodies overnight (Supplementary Table **T2**), incubated with corresponding secondary antibodies (HRP-conjugated anti-mousse, anti-rabbit) for two hours at room temperature, washed and imaged using Odyssey M imager (LICORBio) with ECL Western Blotting Detection Kit (Amersham, RPN2018).

#### Human Angiogenesis & Growth Factor 17-Plex Discovery Assay

The multiplexing analysis was performed by Eve Technologies Corporation (Calgary, Alberta, Canada) using the Luminex® 200™ system (Luminex Corporation/DiaSorin, Saluggia, Italy) with Bio-Plex Manager™ software (Bio-Rad Laboratories Inc., Hercules, California, USA). Seventeen markers were measured in the samples using the Eve Technologies Human Angiogenesis & Growth Factor 17-Plex Discovery Assay Array (HDAGP17) according to the manufacturer’s instructions for use (MILLIPLEX Human Angiogenesis/Growth Factor Magnetic Bead Panel 1, Cat. # HAGP1MAG-12K, MilliporeSigma, Burlington, Massachusetts, USA). The assay sensitivities of the markers range from 0.7 – 8.1 pg/mL. Individual analyte sensitivity values are available in the MilliporeSigma MILLIPLEX® protocol.

#### Statistical analysis

All statistical analyses were performed using GraphPad Prism software (version 10). Each patient was an n=1, and each assay had at least four replicates. Six to eight experimental groups were tested for each patient sample, depending on the number of slices from each patient. Depending on the data distribution, comparison groups, and time points, appropriate statistical tests were employed. P < 0.05 was regarded as statistically significant.

## Acknowledgment

The authors thank Sean Goldfarb, Dr. Klaudia Monika Bednarz, and Dr. Thomas Stroh for providing the Vibratome and Confocal microscope at the Neuro Microscopy Facility; Dr. Peter Metrakos and Dr. Anthoula Lazaris for providing lab space and initial RNA processing at McGill University Health Center; Oscar Boyadjian and Dr. Maryam Tabrizian for NTA equipment and analysis; Dr. Xinyun Su for guiding MSC culture; Transplant Quebec for providing salivary glands. We utilized the computational resources provided by the Digital Research Alliance of Canada for High-Performance Computing. Full text was edited on Grammarly for Grammer correction and language accuracy.

## Research funding

This project is supported by the Canadian Institutes of Health Research (CIHR #249072, #264437) and the Fonds de Research du Québec-Santé Doctoral Training Scholarship (FRQS #317934 https://doi.org/10.69777/317934). The authors declare no conflict of interest.

## Author contributions

A.U.: Conceptualization, methodology, investigations, formal analysis, visualization, data curation, writing and editing, M.T., M.M.: Methodology, review and editing, A.Z., J.P., J.G., H.A.S., N.M.M., M.H., J.W.: Resources and review, S.D.T. Conceptualization, review and editing, project administration and supervision.

## DATA AVAILABILITY

The RNA sequencing data used for the study will be deposited to GeneExpression Omnibus. Correspondence and requests for materials should be addressed to

## Notes

### Competing Interest Statement

The authors have declared no competing interest.

