## Supplementary Tables for "Extracellular Vesicles of Salivary Mesenchymal Stem Cells Mitigate Acute Irradiation Injury: Use of an ex-vivo organotypic human slice tissue culture as a disease model"

Supplementary methods:

**Cells and culture conditions:**

**Labial gland MSC (LMSCs): L**MSCs were isolated and expanded from human minor salivary glands as previously described^11^ and characterized according to the minimal definition criteria suggested by the International Society for Cellular Therapy (ISCT)^35^. The glands were rinsed twice in PBS and their connective tissue was removed gently with forceps and scalpel. Then they were cut into 1-2mm pieces using scalpel no. 10 (Fisher Scientific,36-101-5344) and deposited into 100mm cell culture dish with MEM α with no nucleosides (Gibco, 12561056), along with 1-2% antibiotics and 10% FBS. The pieces gave out cells, which grow to confluency in 10-15 days. To harvest these cells, the explants were gently transferred to another cell culture plate to re-grow, while the released cells were trypsinized (0.05-0.25% trypsin (Gibco 25200-072) for 3-4 min) and centrifuged at 1200rpm (Thermo Electron Corporation IEC Centra GP8R Refrigerated Centrifuge) for 5 min at 4°C). The cell pellet was then resuspended in adequate amount of culture media after counting the cells and seeded for respective experiments**.**

**Submandibular gland (SMG) cells and culture:** The glands were rinsed, and connective tissue removed as previously described. Further, using curved scissors, the gland was minced into a slurry paste. The slurry was then added to a digestion buffer (Liberase TM 1 vial (5mg) in 140 ml of DMEM, with 1% antibodies and 1% glutamine), 35ml buffer for every 2gm of tissue. The tissue was further dissociated using GentleMACS and then incubation at 37°C for 3-5 hours, with vigorous shaking at each half an hour. The digested tissue was then strained with a 70 μm strainer to remove connective tissues and clumps of cells; centrifuged at 900rpm for 5 mins at 4°C. The cells were resuspended in Mammary Epithelial Basal Medium (EpiMax1) (Wisent Inc., 002-010-CL), along with antibiotics and supplements.

**WST-8 proliferation assay:** Human SMG cells were cultured in EpiMax1 cell culture media (Wisent Inc., 002-010-CL). Cells between passages 3-6 were seeded in 96-well cell-culture plates, 3000 cells per well. After 3 days, the cells were exposed to 7.5 Gy IR and treated with different amounts of MSC-CM for two hours. After 24 and 48 hours, cell viability was assessed using Cell counting kit 8 (WST-8, ab228554) according to instructions. 10μl of WST-8 solution was added to each well, incubated for 2 hours followed by measurement of absorbance at 460nm. Cell viability was normalized to the control cells (without IR and without CM treatment). For rate of proliferation, viability 48-hour post-IR was subtracted by 24-hour post-IR.

**SOD activity:** We followed the protocol established previously^79^. Tissue culture media below the trans well inserts was collected at the required time-points, concentrated using Amicon Ultra 0.5ml Centrifugal filters (10kDa cutoff), followed by manufacturer’s instructions for SOD activity assay kit (Invitrogen, EIASODC). As the number of slices could slightly vary from one Trans-well to another, to standardize the assay, we normalized the SOD activity before IR to the SOD activity 24 hours post-IR for each corresponding well.

**Table1:** Patient samples

|  | Male | Females | Total |
| --- | --- | --- | --- |
| Number of subjects | 10 | 8 | 18 |
| Age | 49.7±17.9 | 53.4 ± 26.3 | 51.3 ± 21.4 |

**Table2**: Antibodies

| Antibody | Species | Localization | Dilution | Catalogue No. |
| --- | --- | --- | --- | --- |
| Anti E-Cadherin | Mouse monoclonal | Cell membrane | 1/500 | ab231303 |
| Anti AQP5 | Rabbit polyclonal | Cell membrane | 1/500 | PA5-99403 |
| Anti Alpha-SMA | Mouse monoclonal | Cytoplasm | 1/2000 | Ab7817 |
| Anti Vimentin | Rabbit monoclonal | Cytoplasm | 1/500 | EPR3776 |
| Anti Gamma H2AX | Rabbit monoclonal | Nucleus | 1/500 | Ab81299 |
| Anti-Syntenin (32kDa) | Rabbit monoclonal | EV | 1/1000 | ab133267 |
| Anti-Calnexin(68kDa) | Rabbit monoclonal | Endoplasmic Reticulum | 1/1000 | ab133615 |
| Anti-Human CD63 | Mouse monoclonal | EV | 1/1000 | BD 556019 |
| Anti TSG-101 | Rabbit monoclonal | Endosome sorting complex | 1/1000 | EPR7130(B) |

**Table3**: qPCR primers:

|  | Forward | Reverse |
| --- | --- | --- |
| *gH2AX* | 5'-CGGGCGTCTGTTCTAGTGTT-3' | 5'-GGTGTACACGGCCCACTG-3 |
| *FDXR* | 5'-CACCCCGAGTAGGAGCAG-3' | 5'-GGAGAAATGGTGGCAGAAG-3' |
| *MDM2* | 5'-TGGTGAACGACAAAGAAAACG-3' | 5'-GTAACTTGATATACACCAGCATCAA |
| *P21/CDKN1A* | 5'-ACTCTCAGGGTCGAAAACGG-3' | 5'-GATGTAGAGCGGGCCTTTGA-3' |

Supplementary data:

**Table 4**:

|  | *Gene* | logFC | p-adj | Slice(avg) | Gland(avg) |
| --- | --- | --- | --- | --- | --- |
| Ductal | *KRT5* | 0.00775364 | 0.98726921 | 13.0693636 | 13.1294137 |
|  | *KRT7* | 2.35599223 | 2.26E-12 | 17.4187098 | 15.8716429 |
|  | *KRT14* | 0.12335072 | 0.81903209 | 13.1844578 | 13.1220374 |
|  | *KRT19* | 2.13936858 | 1.17E-10 | 15.990292 | 14.5703501 |
|  | *EPCAM* | 0.74618302 | 3.07E-05 | 14.1501032 | 13.6409392 |
|  | *CDH1* | 0.47082167 | 0.03009916 | 14.2713492 | 13.9393446 |
|  | *TJP1* | 0.8523322 | 8.30E-05 | 12.6079394 | 12.0327609 |
|  | *VIPR1* | -0.8686697 | 0.00087931 | 9.3614634 | 9.95247165 |
|  | *EPGN* | 9.94117472 | 1.81E-15 | 10.3499393 | 6.0436113 |
| Acinar | *AQP5* | -4.8604854 | 1.63E-22 | 12.2263818 | 15.6109448 |
|  | *CHRM3* | -3.62816 | 5.56E-20 | 10.5086802 | 12.9634235 |
|  | *BHLHA15* | -6.6668195 | 2.27E-53 | 9.82548452 | 14.4460429 |
|  | *SLC12A2* | -1.7186483 | 3.60E-06 | 12.4893552 | 13.6364718 |
|  | *MUC19* | -6.6266793 | 6.65E-13 | 6.71725934 | 10.954246 |
|  | *AMY1a* | -4.8672825 | 1.53E-06 | 15.5980687 | 19.3350417 |
|  | *ASCL2* | -2.4866777 | 2.46E-17 | 8.60925221 | 10.2716423 |
|  | *ASCL3* | -8.0682722 | 1.87E-13 | 3.28330363 | 5.37839438 |
| Myoepithelial | *TRPV4* | -3.1486805 | 8.87E-14 | 8.96468555 | 11.2238474 |
|  | *ACTA2* | -6.4716519 | 3.21E-93 | 8.30338208 | 12.814555 |
|  | *CNN1* | -4.6454333 | 6.77E-73 | 7.57219493 | 10.7356932 |
| Proliferation | *PCNA* | 1.40221788 | 3.81E-07 | 11.5735932 | 10.6309829 |
|  | *KI67* | 1.14664443 | 0.23305409 | 6.79769906 | 6.26915372 |
|  | *MYC* | 2.3462065 | 2.56E-10 | 13.2308031 | 11.5584507 |
| Macrophage | *LY6C* | NA |  |  |  |
|  | *ADGRE1 (F4/80)* | NA |  |  |  |
|  | *CD68* | 2.10967025 | 1.11E-06 | 10.917813 | 9.4736109 |
| M1 | *CD14* | -0.4114212 | 0.48572485 | 12.1904628 | 12.5447909 |
|  | *CD80* | NA |  |  |  |
|  | *CD86* | -0.2238081 | 0.83281427 | 6.50216827 | 6.78105693 |
|  | *CD16* | Not found |  |  |  |
| M2 | *CD163* | -0.2631377 | 0.69884234 | 8.54589654 | 8.70488982 |
|  | *CD206* | NA |  |  |  |
|  | *CD209* | NA |  |  |  |
|  | *CCL2* | 4.40990828 | 4.94E-08 | 13.4590478 | 10.445698 |
| Endothelial cells | *CD31/PECAM1* | -1.2137985 | 0.0638188 | 8.82266566 | 9.73186974 |
