## Supplementary figures and images for "Extracellular Vesicles of Salivary Mesenchymal Stem Cells Mitigate Acute Irradiation Injury: Use of an ex-vivo organotypic human slice tissue culture as a disease model"

### Supplementary Figure 2

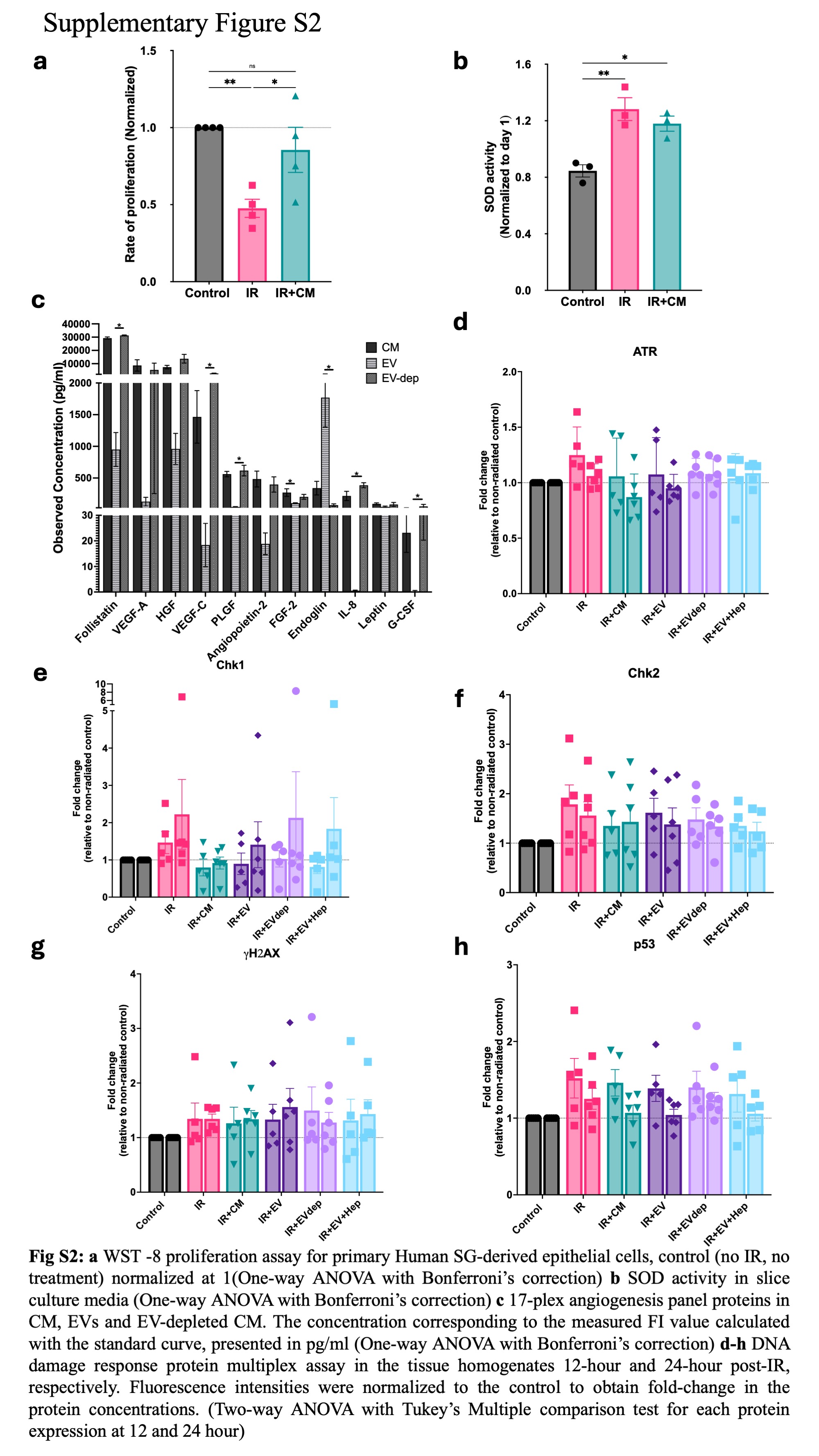
