## Supplementary Figure 1 for "Extracellular Vesicles of Salivary Mesenchymal Stem Cells Mitigate Acute Irradiation Injury: Use of an ex-vivo organotypic human slice tissue culture as a disease model"

### Supplementary Figure S1

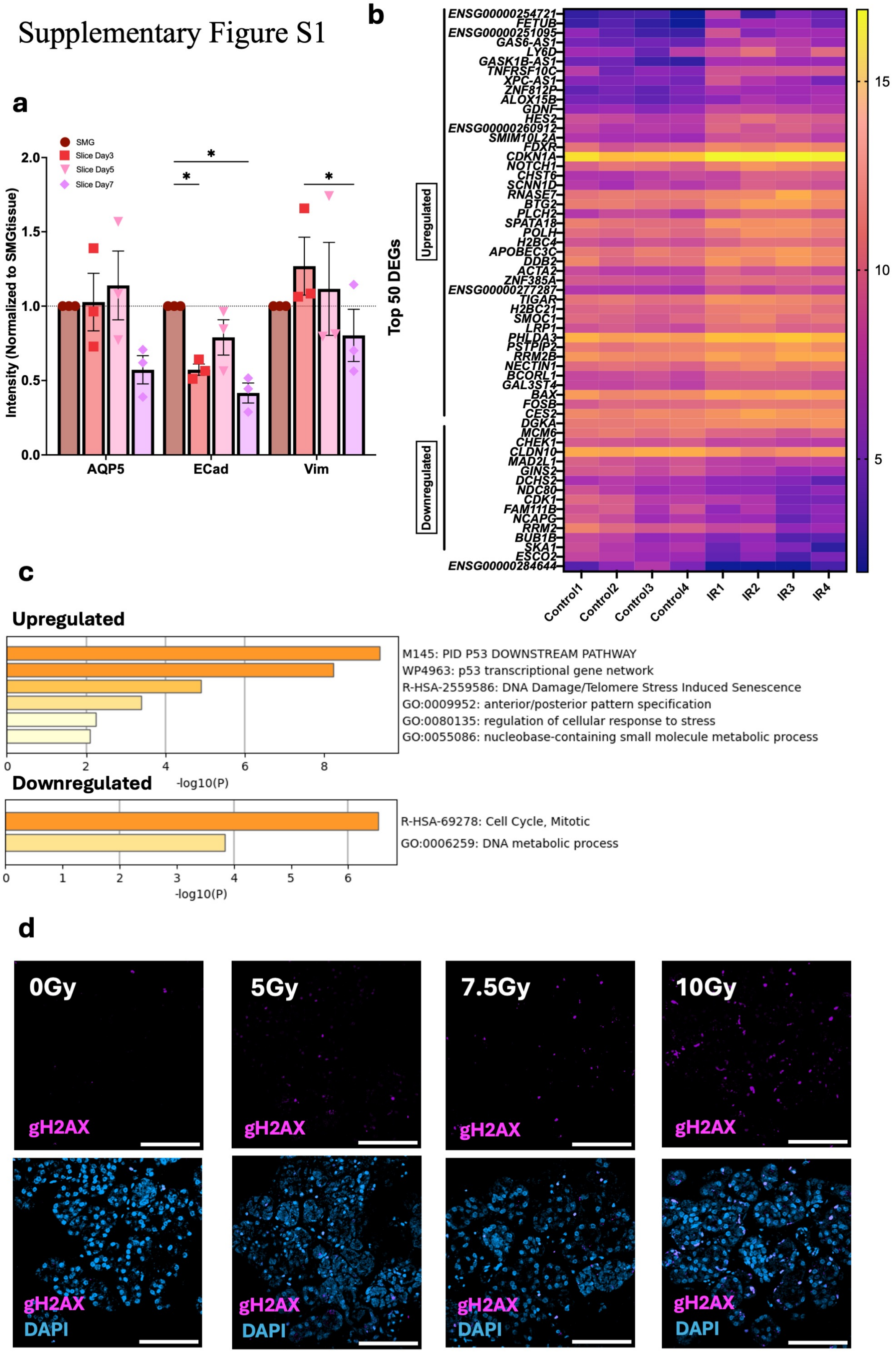


**Fig S1. a** Intensity analysis for AQP5, ECadherin and Vimentin in intact SMG, slices at day 3, 5 and 7(Normalized to intact SMG, Two–way ANOVA, with Tukey's multiple comparison test). **b** Heatmap of top 50 genes expressed in Control and IR slices from bulk RNA-seq analysis. The colour scale represents DeSeq2 normalized expression values for each sample. GSEA: KEGG database, FDR cut-off=0.1 **c** Functional pathways identified through Metascape for upregulated and downregulated genes **d** Immunofluorescence staining for gH2AX in response to various IR doses.
